# Water position prediction with SE(3)-Graph Neural Network

**DOI:** 10.1101/2024.03.25.586555

**Authors:** Sangwoo Park

## Abstract

Most protein molecules exist in a water medium and interact with numerous water molecules. Consideration of interactions between protein molecules and water molecules is essential to understanding the functions of the protein. In computational studies on protein functions, either implicit solvation or explicit solvation methods are used to consider the effect of water on the protein. Implicit solvation methods consider water as a continuous solvent and have lower computational costs than explicit methods that consider water as a collection of individual water molecules. However, some water molecules have specific interactions with protein molecules, which are critical to protein function and require explicit treatment to consider these specific interactions. Thus, as a compromise between computational cost and consideration of specific interactions, hybrid methods use explicit consideration of water molecules with specific interaction with protein molecules while considering other water molecules implicitly. Prediction of the water positions having specific interaction is required to perform such hybrid methods, where various water position prediction methods have been developed. However, currently developed water position prediction methods still require considerable computational cost. Here, we present a water position prediction method with low computational cost and state-of-the-art prediction performance by utilizing SE(3)-an equivariant graph neural network. The introduction of a graph neural network enabled the consideration of the atom as a single data point, which makes computational costs less than our previous water prediction method using a convolutional neural network, which considers an atom as multiple data points. Our new water position prediction method, WatGNN, showed an average computation time of 1.86 seconds while maintaining state-of-the-art prediction performance. The source code of this water prediction method is freely available at https://github.com/shadow1229/WatGNN.

## 1. Introduction

Many protein molecules exist in an aqueous solution and have interactions between protein molecules and water molecules. These interactions between protein molecules and water molecules can affect the structure and function of proteins^1–13^. In computational studies of protein structure and function, implicit solvation and explicit solvation methods have been used to consider water molecules of protein molecules. The implicit solvation method considers water as a continuous solvent medium, which enables the solvation effect to be considered with relatively low computational costs^14^, which makes the implicit solvation method favorable for protein docking calculations that require lots of calculations^15–17^. However, implicit solvation cannot correctly consider specific atomic interactions between water molecules and protein atoms, including hydrogen bonds between water and protein molecules^18^. Since hydrogen bonds between protein molecules and water molecules affect proteins’ surface structure and function^19, 20^, the consideration of explicit water molecules in protein docking methods has been emphasized^21, 22^. To consider specific interactions between protein molecules and water molecules with low computational cost, hybrid methods have been proposed explicitly considering water molecules with specific interactions with protein molecules and implicitly considering the rest^23–25^.

To perform such hybrid methods, the positions of water molecules that have specific interactions with protein molecules must be predicted. Water position prediction methods developed so far include methods based on explicit-water molecular dynamics (MD) simulations^26–33^, methods based on integral equation theory^34–38^, methods based on hydrogen bond geometry between protein molecules and water molecules^39–42^, methods based on statistical potential between protein molecules and water molecules^43, 44^, and methods based on Convolutional Neural Network (CNN)^45, 46^.

With the development of deep learning, there are water position prediction methods using deep-learning-based methodology, including the CNN-based methods mentioned above. These methods have shown high prediction performance via high capability in structure pattern recognition of CNN models^45, 46^. However, the traditional CNN-based model predicts water positions on grid points that span the input protein structure. Thus, to achieve water position prediction with sub-angstrom accuracy, the spacing between grid points should be shorter than 1 Å, producing convolutional calculations for tensors much larger than the number of atoms in the protein structure given as input and causing considerable computational cost. Additionally, previous CNN-based water prediction methods would make different results from the same input structure with different orientations because the traditional CNN model cannot ensure equivariance with the rotation of the input.

In this research, we present the fast water position prediction method WatGNN, which uses SE(3)-equivariant Graph Neural Network^47, 48^ (SE(3)-GNN) to reduce computational cost and ensure equivariance of rotation and translation of the input. The neural network processes input protein structure as a graph, where each atom in the protein is regarded as a node. Node features are updated in the translation-equivariant and rotation-equivariant manner by using the relative position of nodes for the generation of messages between nodes, where the harmonics-based kernel was used for processing the orientation term of the relative position to preserve rotational equivariance.

The water position prediction result of WatGNN are compared to previous methods with various approaches: CNN-based method GalaxyWater-CNN^45^, statistical potential method GalaxyWater-wKGB^43^, integral theory method 3D-RISM^38^, and a geometry-based method FoldX^49^. When compared to a high-resolution crystal set, the accuracy of water position prediction using WatGNN is substantially better than the accuracy of the prediction using GalaxyWater-CNN, which has shown the best performance among previous methods^45^, while the average computational cost for prediction of water position on the set was 1.86 seconds on i9-12900k CPU and RTX-4090 GPU, which is 12.8 times faster than GalaxyWater-CNN.

## 2. Methods

### 2.1 Overview

WatGNN predicts water positions from a given protein structure with SE(3)-equivariant Graph Neural Network model, as shown in **Figure 1**. The network model takes the atom types and positions of atoms in the structure of the input protein molecule and returns the predicted positions of water molecules and probability-like scores of each predicted position. To predict water positions with specific atomic interactions, such as hydrogen bonds with protein molecules, WatGNN predicts water positions near non-carbon atoms(=N, O, S, P, halogens, and metal) of input protein and compound molecules, which atoms could make interaction such as hydrogen bonds or ionic interactions with water molecules. A probe node was introduced to predict multiple water molecule positions near non-carbon atoms of input molecules. Probe nodes are artificial nodes with four nodes placed near each non-carbon atom and are used to predict water positions. Each probe node can predict up to 2 positions of water molecules, where the first predicted position is a predicted water position with a hydrogen bond with protein directly, and the other position is a predicted water position with a hydrogen bond only with other water molecules. From the predicted positions and scores on the probe nodes, the positions are sorted by scores and placed sequentially without having clashes between predicted positions by clustering predicted positions.

**Figure 1.**
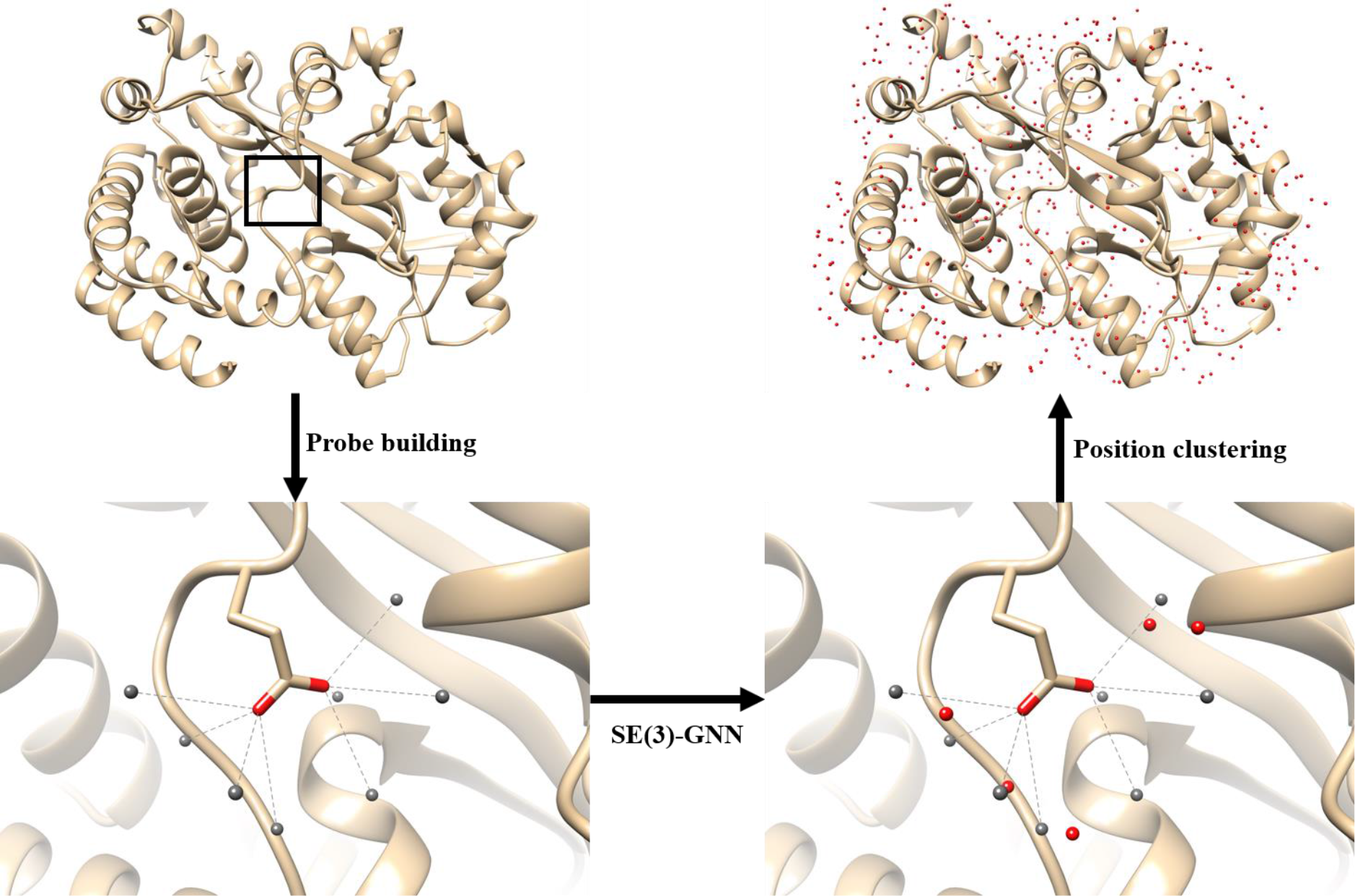
WatGNN predicts the positions of water molecules near the given structure of a protein by SE(3) - equivariant Graph Neural Network. WatGNN predicts up to two positions for each probe node placed near a non-carbon atom in the input structure. In the figure, probe nodes are shown as gray spheres, and predicted water positions are shown as red spheres.

### 2.2 Workflow of WatGNN

The input of WatGNN is the atom type and positions of atoms in the given protein structure. The input structure is processed into a bidirectional graph, where a node represents atoms and probes of the protein molecule, and edges are established between i) intra-residual atom pairs that share the same protein residue, ii) non-carbon atom pairs whose pair length is shorter than 10Å, iii) pairs of a non-carbon atom and its probe. For the node feature of the graph, the atom type of the node was used, where ten atom types were considered, corresponding to C, N, O, S, P, halogens, alkali metals, non-alkali metals, probe, and other atoms. The edge feature has three features of edge types: intra-residual connection, non-carbon atom pair connection, and probe – non-carbon atom connection, as shown in **Figure 2**.

**Figure 2.**
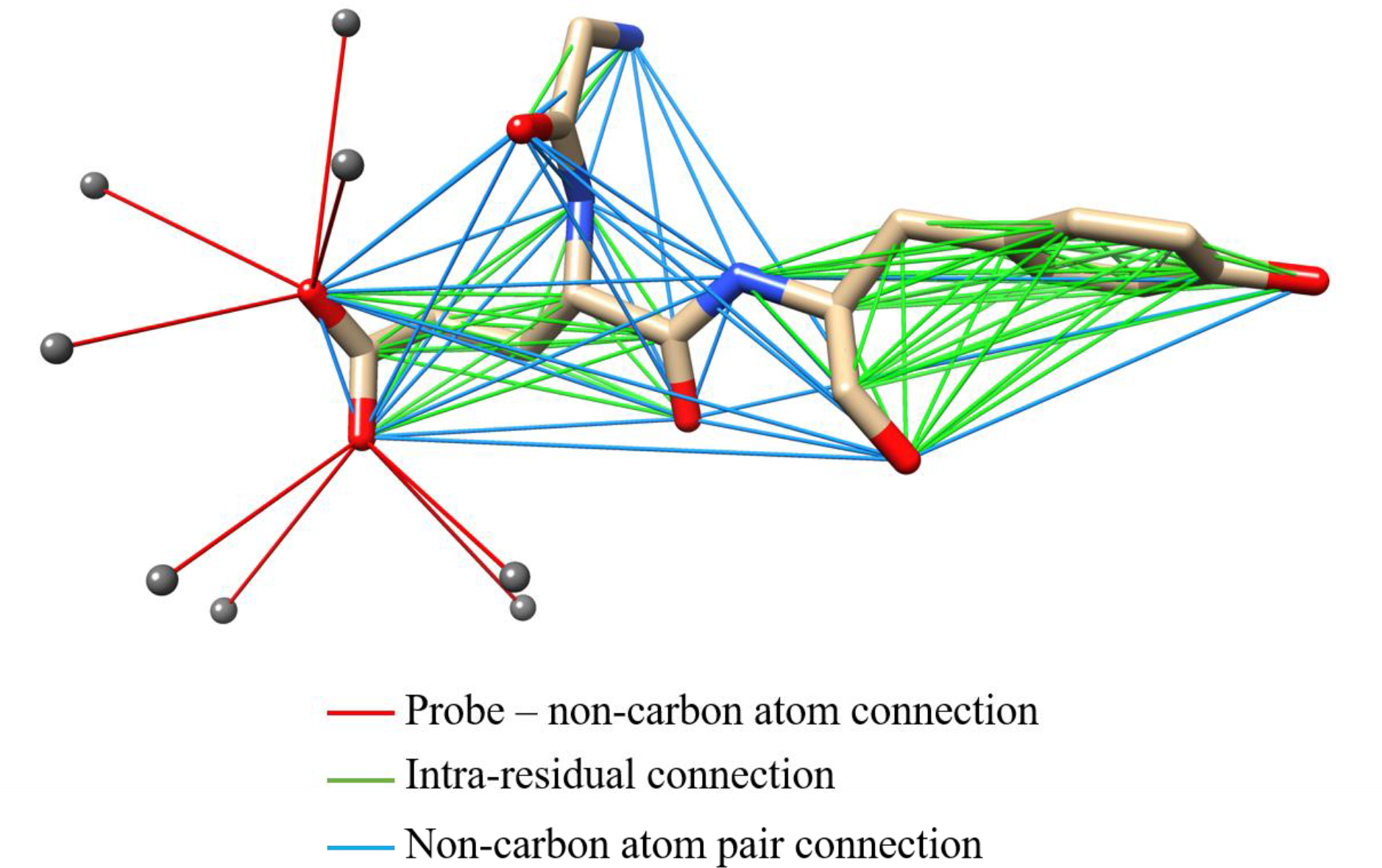
This figure shows an example of edges and their types in WatGNN. Gray spheres represent the probe nodes. For the convenience of visualization, probe nodes for oxygen atoms in carboxyl groups are shown only.

For the placement of probe nodes, probe nodes are placed based on the local dipole orientation vector of the non-carbon atom (*d̂*) and center orientation vector of the non-carbon atom (*ĉ*) for the conservation of geometric properties of hydrogen bond between protein atoms and water molecule as shown in **Figure 3(a)**. Inspired by our previous research^43^, the local dipole orientation vector is defined as orientation vector of the local dipole vector which defined as *r*_*p*_ − ⟨*r*_*q*_⟩, where *r*_*p*_ is the coordinate of the non-carbon atom *p* that probe node is placed, and ⟨*r*_*q*_⟩ is the average coordinate of {*q*} non-hydrogen atoms that are connected to atom *p* by chemical bond,. The center orientation vector is defined as the orientation vector of the coordinate of the non-carbon atom from its average coordinate of non-hydrogen atoms in its residue. From the two orientation vectors, three axis vectors for placement of the probe are calculated by Equation (1), where 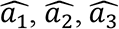 is three axis vectors. Additionally, if *ĉ* and *d̂* is parallel, random unit vector perpendicular to 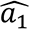 was used as 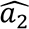.

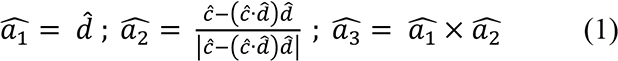

**Figure 3.**
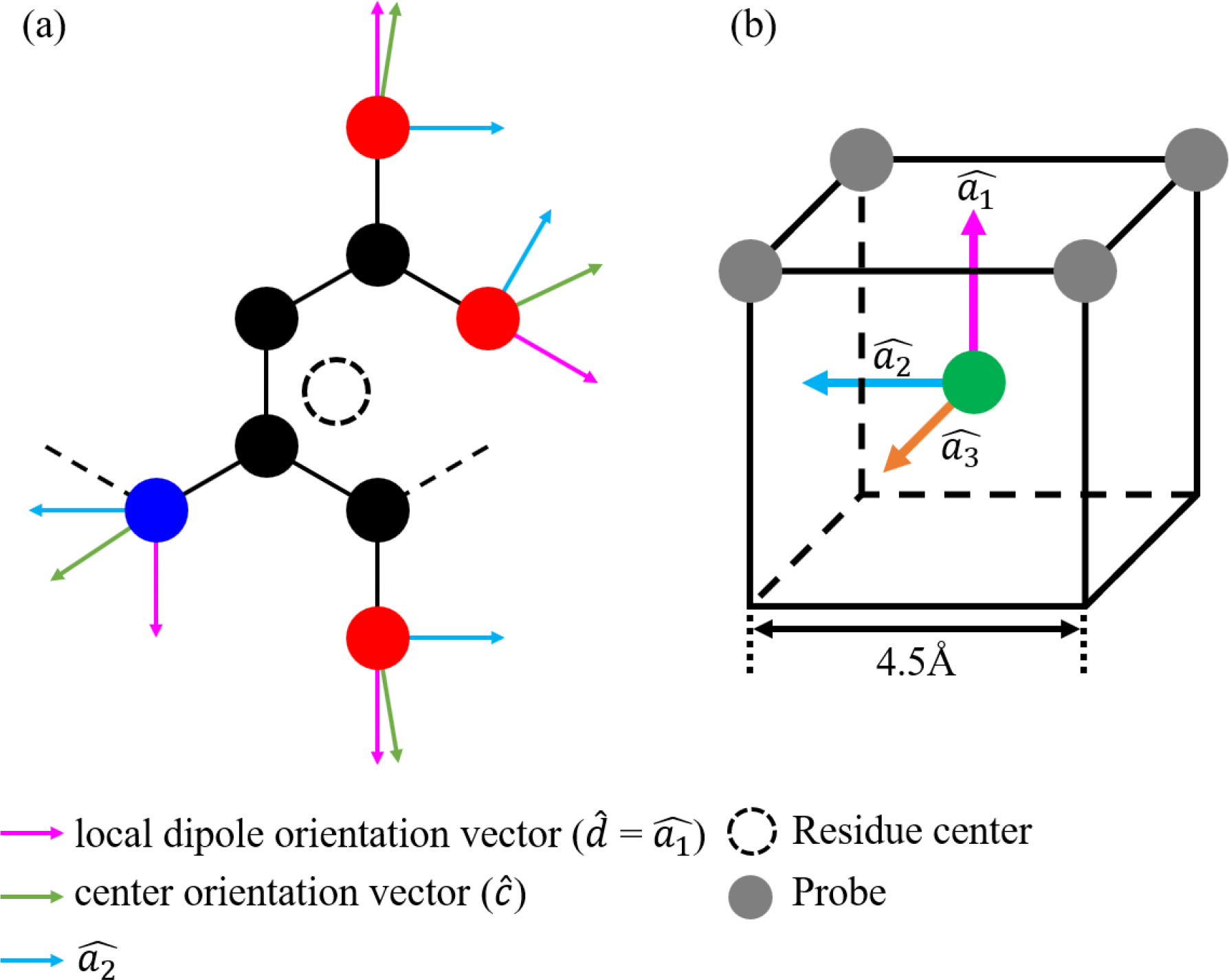
Probe placement for non-carbon atoms in WatGNN. (a) shows orientation vectors for the input structure, (b) the location of probes for a given non-carbon atom, and three axis vectors for probe placement. In figure (b), non-carbon atom is located the center of 4.5 Å cube, which axis is parallel to 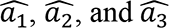. Four probes are located at the vertices of the cube.

4 probe nodes are placed as **Figure 3(b)** and Equation (2), where *r*_*prb,2j+k*_ is the coordinate of 2*j*+*k* th probe.

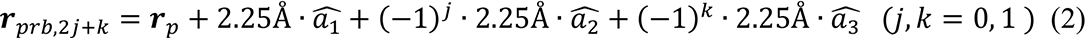

The network structure of WatGNN is shown in **Figure 4**. WatGNN was created with reference to cg2all^47^, which consists of encoding, interaction, and structure modules. An encoding module was used to encode node features of 10 atom types into 16 scalar features. Node and edge features are processed with an interaction module, which performs interaction between nodes with 6 SE(3)-Transformer blocks^50^ followed by a ConvSE3^50^ operation. SE(3)- Transformer blocks consist of a sequence of SE(3)-Transformer, layer normalization(LayerNorm)^51^, and exponential linear unit(ELU) activation^52^, where SE(3)-

**Figure 4.**
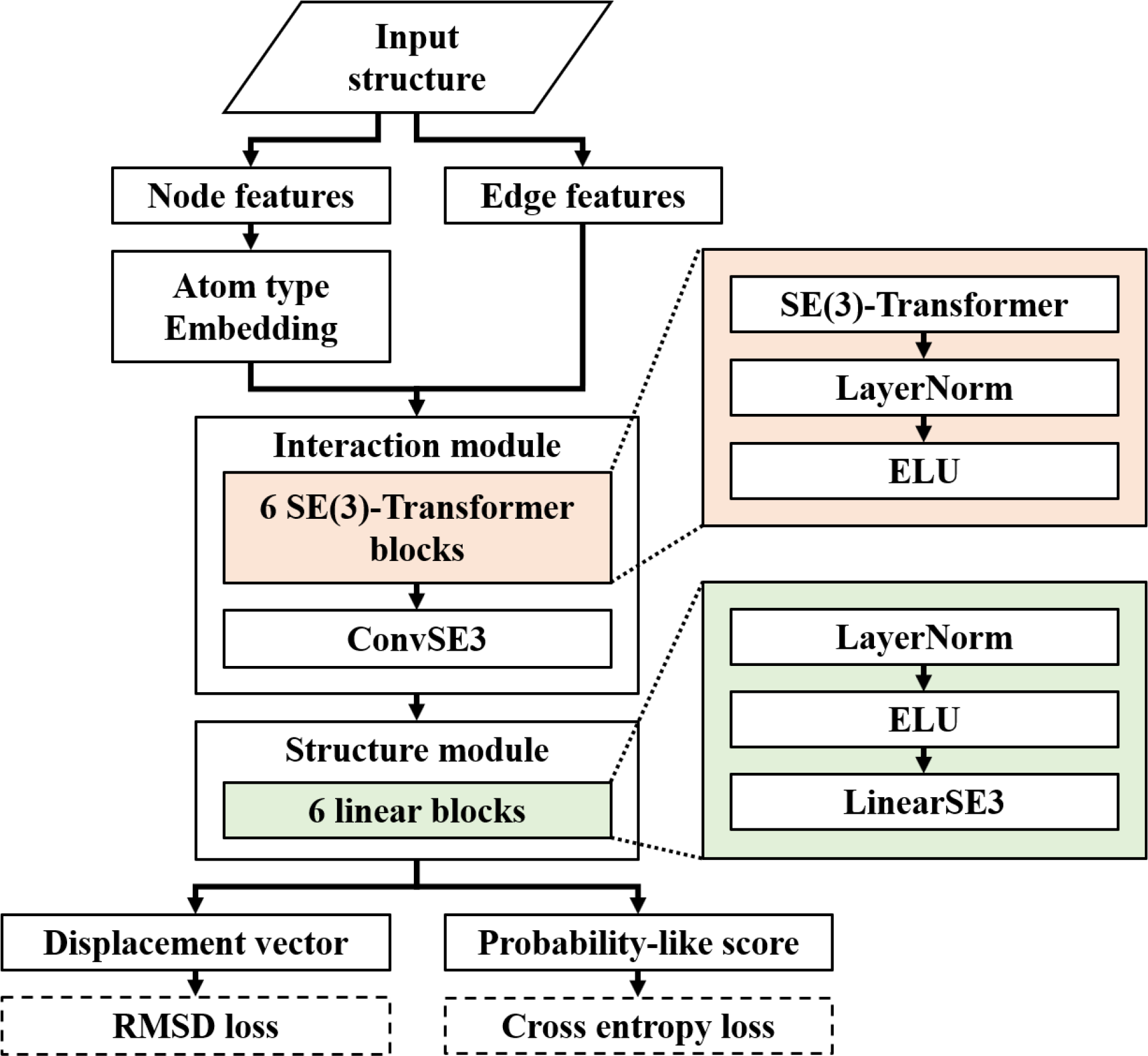
The network structure of WatGNN with its loss functions. In the figure, SE(3)- transformer block and linear block are colored, and the components of the blocks are shown to be the same color as the block.

Transformer performs 8-head self-attention between a node and its neighboring nodes that connected by an edge, and ConvSE3 performs Graph convolution. After SE(3)-Transformer, node features are processed into 64 scalars, 64 vectors, and 64 traceless matrices, and ConvSE3 transforms node features into 64 scalars and 32 vectors. Finally, the structure module converts node features into two scalars and two vectors via six linear blocks, where linear blocks consist of a sequence of LayerNorm, ELU, and LinearSE3^50^, where LinearSE3 performs a 1x1 convolution-like operation on each node. The resulting node features on probe nodes are trained to predict the probability of neighboring water molecules near the probe and the displacement vector from the probe to the predicted position. Additionally, to predict both water positions that have hydrogen bonds with protein atoms and have hydrogen bonds with other water molecules, the first channel of each scalar and vector feature is trained to predict probability and displacement for water molecules that have a hydrogen bond with protein atoms, and second channels are trained to predict probability and displacement for water molecule that having a hydrogen bond with other water molecules only.

After generating predicted positions, position clustering was performed to place predicted water positions without clash. To cluster positions, predicted water positions and their probability from the neural network are sorted with probability scores in descending order. Then, predicted water positions are placed only if the newly placed water position does not have a neighboring previously placed water position within 2Å.

### 2.3 Network training

A high-resolution X-ray crystal structure database was constructed to train and evaluate the network without redundancy. The database was gathered from Protein Data Bank (PDB) and built with PISCES protein sequence culling server^53^ on September 21, 2023, with resolution better than 2.0Å, R-free values lower than 0.25, sequence identity lower than 30%, and maximum number of non-hydrogen protein atoms are limited to 5000. Also, crystal structures have a number of well-resolved water molecules, defined as B-factor <40 between 5 and 20% of the number of protein atoms. The total number of protein structures was 7971, of which 3971 were used as a training set, 300 were used as a validation set, and the remaining 3700 structures were used as a test set.

For training and evaluation, atoms from protein and other compound molecules without water molecules were used as inputs, and well-resolved water molecules defined as B-factor <40 were used for ground truth. In the training, the ground truth of the existence of the water position near the probe and the displacement of the water position from the probe position were assigned to each probe. To train the first channel for prediction of water position having protein-water hydrogen bond, ground truth of the first channel of the probe is assigned to the water position that exists inside of the probe’s covering cube and having a hydrogen bond with non-carbon atom, which defined as having neighboring non-carbon atom within 3.5Å and having 100° or larger angle that made between the position of water, the non-carbon atom, and the average coordinate of non-hydrogen atoms forming chemical bonds to the non-carbon atom. The probe’s covering cube has 4.5Å of side length, shares a non-carbon atom as a vertex, and has the probe at the center. To train the first channel for prediction of water position having water-water hydrogen bond, the ground truth of the second channel is assigned to the water position that exists inside of the probe’s covering cube and having other neighboring water position within 3.5Å and having no hydrogen bond with any non-carbon atom which criteria is described above. If two or more positions are assigned to the same probe, the water position closest to the non-carbon atom is assigned to the ground truth. To train the network without severe overtraining, a batch size of 4 randomly truncated protein structures was used as an input, which truncated a sphere radius of 15Å and has a center on the position of a random protein atom. For the Loss function, weighted cross entropy with a weight of 4 for having water was used for scalar output, which predicts a probability-like score for the water position and RMSD between predicted displacements and displacements from the positions of ground truth and the positions of probes was used for vector output, which predicts displacement between predicted water position and position of the probe. The total loss function is the sum of weighted cross entropy and RMSD. Training the network was done with Pytorch^54^, with 150 epochs of training using Adam optimizer with a learning rate of 0.001 and exponential decaying rate of 0.99.

### 2.4 Performance evaluation

Coverage of well-resolved crystallographic water positions defined as crystallographic water positions having B-factor <40 and RMSD of predicted water position from well-resolved crystallographic water positions was used to evaluate Water position prediction performance. The coverage was defined as the fraction of well-resolved crystallographic water positions within a specific distance cutoff from the predicted positions among all well-resolved crystallographic water positions. This research measured coverage with distance cutoffs of 0.5Å, 1Å, 1.5Å was measured. The coverage and RMSD were calculated after matching at most one crystallographic water position to each predicted position and were calculated for a different number of predicted positions, ***N***_pred_, which was set to ***N***_pred_ = *n****N***_cryst_, where ***N***_cryst_ is a number of well-resolved water positions in the crystal structure, varying *n* from 1 to 10.

The performance of WatGNN was compared with GalaxyWater-CNN^45^, GalaxyWater- wKGB^43^, 3D-RISM^55, 56^, and FoldX^49^ with a single-protein comparison set and protein-compound comparison set, which was used as benchmark sets in a previous study, GalaxyWater-CNN^45^. The single protein comparison set consists of 92 high-resolution X-ray crystal structures with a resolution better than 1Å and was released before March 14th, 2015. and does not overlap with the training set. For the protein-compound comparison set, 397 structures were curated from the PDBBind set^57^, which contains high-resolution X-ray crystal structures. For curation, resolution better than 2Å, sequence identity lower than 30%, Tanimoto similarity of compounds less than 50%, and crystal structures that have a number of well-resolved water molecules, which is defined as B-factor <40 between 5 and 20% of the number of protein atoms was used. The training, validation, and test sets lists are provided at https://github.com/shadow1229/WatGNN.

### 2.5 Running other prediction methods for comparison

For performance comparison, predictions from GalaxyWater-CNN^45^, GalaxyWater- wKGB^43^, 3D-RISM^55, 56^, and FoldX^49^ were compared with the predictions from WatGNN. Predictions with GalaxyWater-wKGB, 3D-RISM, and FoldX followed the same method as the previous research, GalaxyWater-CNN.

For detailed information on other methods, GalaxyWater-wKGB was run on the web server (http://galaxy.seoklab.org/wkgb). It does not require any options and predicts water positions with their corresponding scores. Prediction with 3D-RISM was performed by using the “rism.snglpnt” program from the AmberTools simulation suite^58^ to generate a water density map and locate the water position from the map by “Placevent” ^59^ from AmberTools.

The SPC/E water model and ff99SB and General AMBER Force Field (GAFF) were used for the force field. For the charges of the non-protein compound, the AM1-BCC charge was used with protonation states given in the PDBBind refined set. Also, 55.5M water with 0.005 M sodium and chloride ions was used as a solvent for 3D-RISM. For 3D-RISM, Kovalenko-Hirata (KH) approximation^60^ was used to close the integral equation, with Grid spacings of 0.5 Å and a minimum buffer of 14 Å between the solute and the grid box.

FoldX was run using the FoldX CrystalWaters program^49^, with the default options (‘lowAffinityMetal’ of -3 and ‘numWaterPartners’ of 2).

For computation time comparison, GalaxyWater-CNN, 3D-RISM, and FoldX were compared with WatGNN, which performed on an Intel i9-12900k CPU and Nvidia RTX 4090 GPU.

## 3. Results

### 3.1 Result of network training

Figure 5 plots the coverage and RMSD of WatGNN for the training, validation, and test sets, which shows minimal overtraining on the training set. As an example, water molecules were predicted with an average coverage of 52%, 50%, and 50% when ***N***_pred_/***N***_cryst_ = 1 and 71%, 68%, and 67% when ***N***_pred_/***N***_cryst_ = 3 for the training, validation, and test sets, respectively.

**Figure 5.**
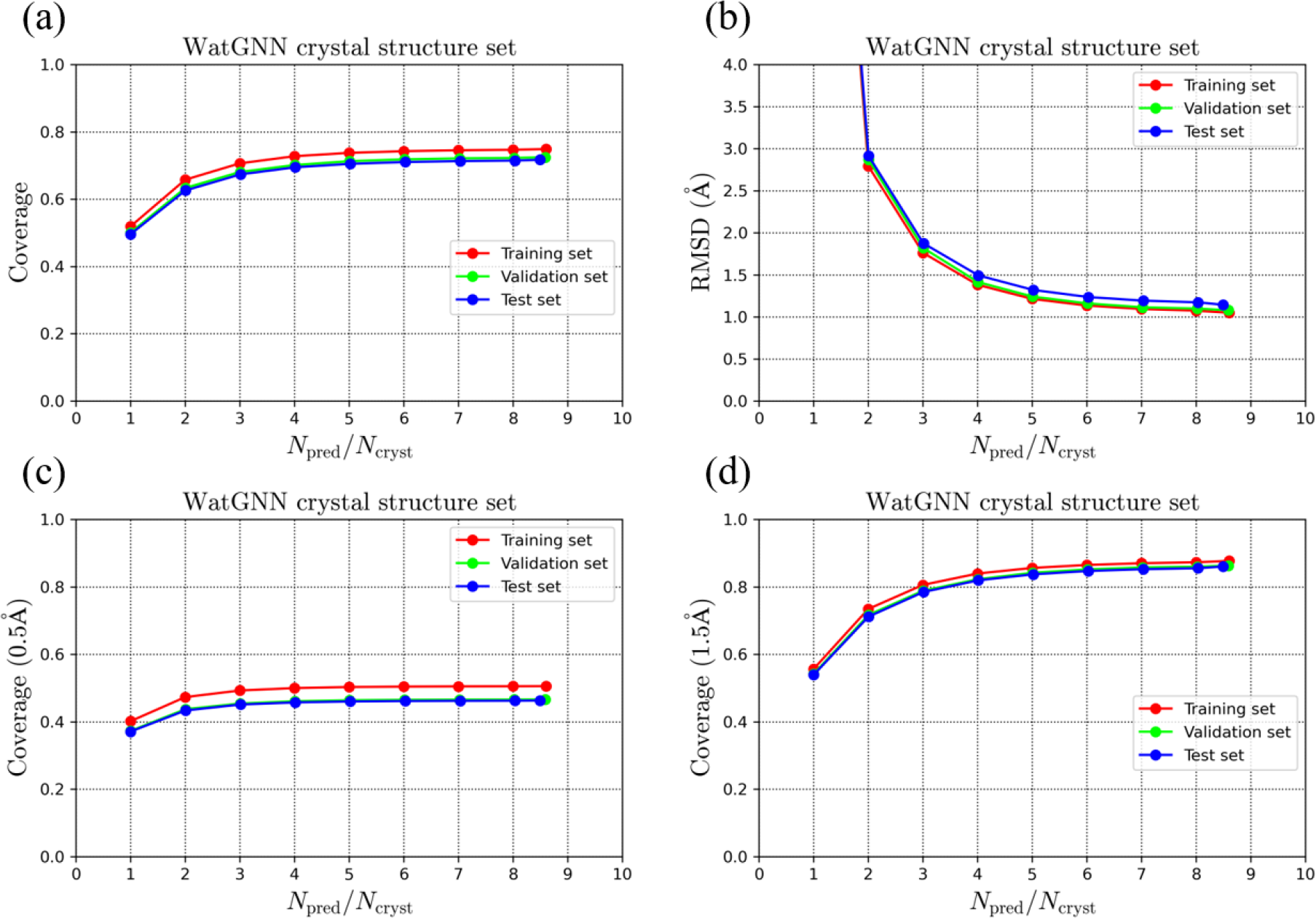
The prediction performance of WatGNN on its training, validation, and test set: a), c), d) Average coverage, and d) Average RMSD of the predicted water molecules vs. the number of predicted water positions scaled by the number of water positions in the crystal structure was shown. For the average coverage, distance cutoff criteria of a) 1.0 Å, c) 0.5 Å, and d) 1.5 Å were used, respectively.

### 3.2 Results on the single protein comparison set

The water position prediction performance and computation time of WatGNN were compared with GalaxyWater-CNN, GalaxyWater-wKGB, 3D-RISM, and FoldX on the single-protein comparison set, which is independent of WatGNN training set and has been used for performance comparison in previous studies^43, 45^. As can be seen from the comparison shown in Figure 6, the average prediction performance of WatGNN is better than other methods used to predict a small number of positions. For example, WatGNN shows an average coverage with 1Å criteria of 54% and 76% when ***N***_pred_/***N***_cryst_= 1 and 3 respectively, while GalaxyWater-CNN shows coverage of 45% and 75%. Also, WatGNN has shown better performance for more strict criteria, where WatGNN shows an average coverage with 0.5Å criteria of 45% and 58% when ***N***_pred_/***N***_cryst_= 1 and 3 respectively, while GalaxyWater-CNN shows coverage of 30% and 47%. Also, the average computation time of WatGNN was less than other methods, where the average computation time for WatGNN was 1.86 s, compared with 23.77s, 398.77s, and 5.72s for GalaxyWater-CNN, 3D-RISM, and FoldX, respectively.

**Figure 6.**
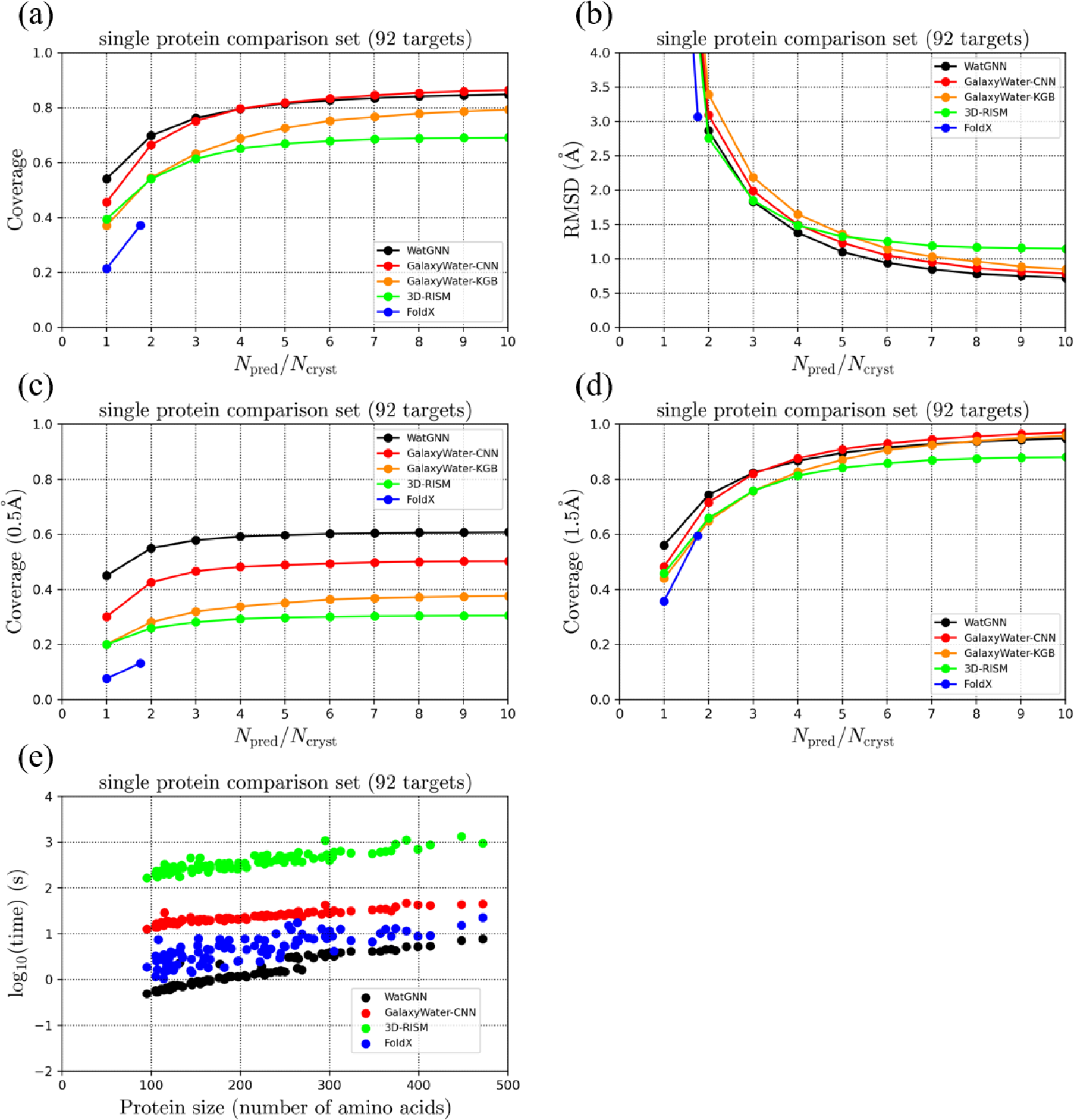
The prediction performance comparison of WatGNN on a single protein comparison set: a), c), d) Average coverage, and d) Average RMSD of the predicted water molecules vs. the number of predicted water positions scaled by the number of water positions in the crystal structure was shown. For the average coverage, distance cutoff criteria of a) 1.0 Å, c) 0.5 Å, and d) 1.5 Å were used, respectively. e) shows computation time vs. the number of amino acids in the input protein structures.

As an example of prediction performance, the water sites predicted with WatGNN and GalaxyWater-CNN are compared in Figure 7. The figure shows that predicted positions with WatGNN are closer to water sites in the crystal structure than those with GalaxyWater-CNN. This demonstrates the difference between WatGNN and GalaxyWater-CNN, where WatGNN predicts the displacement vector between the probe node and water position. At the same time, GalaxyWater-CNN generates a score map on the grid points and selects grid points with a high score to predict water positions, which limits the precision of the predicted position by the spacing of the grid. Furthermore, the computation time for grid-based methods such as GalaxyWater-CNN, GalaxyWater-wKGB, and 3D-RISM could be longer than WatGNN due to score calculations performed at every grid point that covers the input protein structure, which is far more than the number of probes in WatGNN. However, the coverage of predicted positions of WatGNN could be worse than grid-based methods when numerous positions are predicted due to the limitation of WatGNN that predicts positions within 4.5 Å of probe nodes. However, the grid-based model does not have such limitations.

**Figure 7.**
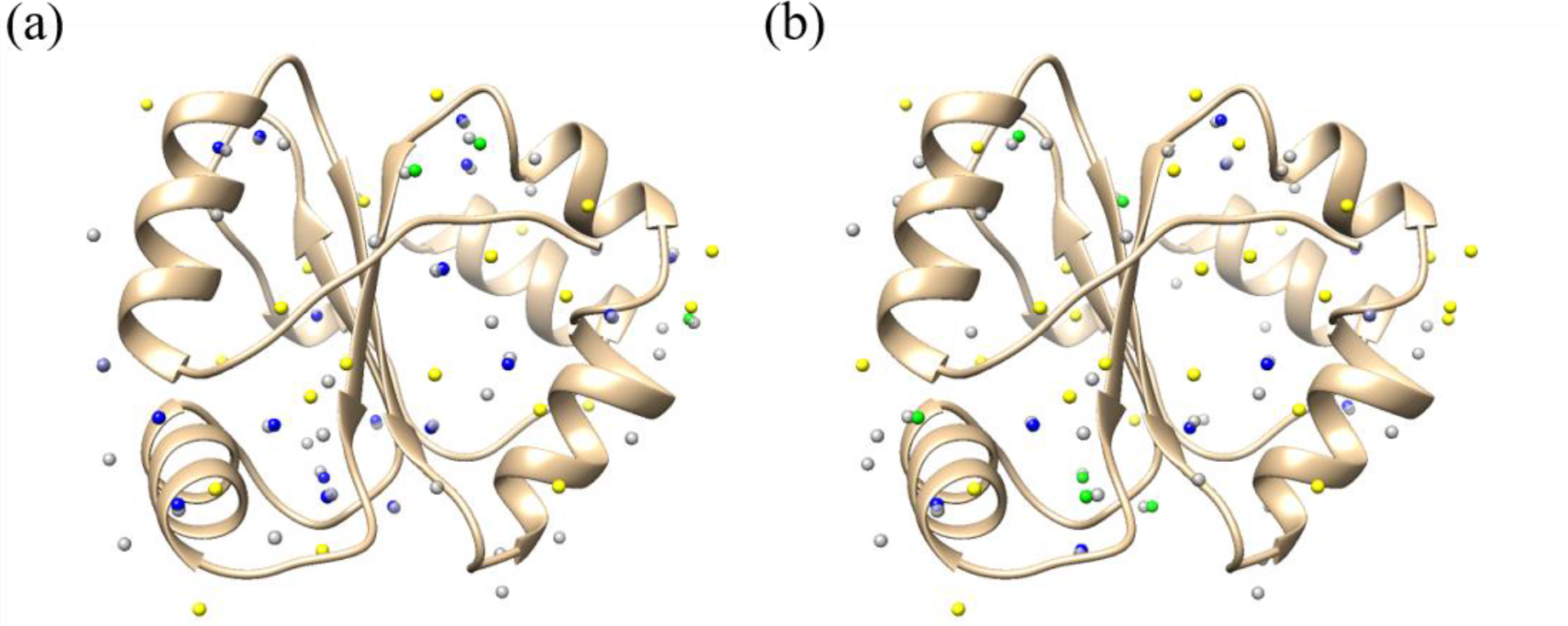
An example case of water site prediction (PDB ID: 2FWH) done with (a) WatGNN and (b) GalaxyWater-CNN, with 47 positions predicted, which is the number of crystallographic water molecules. Gray spheres represent predicted water positions, and colored spheres (yellow, green, and blue) represent crystallographic water sites. Blue spheres indicate crystallographic water molecules with corresponding predicted water sites within 0.5 Å, and green spheres indicate crystallographic water molecules with corresponding predicted water sites within 1 Å, and yellow spheres indicate crystallographic water molecules for which no predicted water site is found within 1 Å. In this figure, water position prediction with WatGNN showed coverage of 47% (22 out of 47) for 1 Å criteria and 40% (19 out of 47) for 0.5 Å criteria. In comparison, GalaxyWater-CNN showed coverage of 34% (16 out of 47) for 1 Å criteria and 21% (10 out of 47) for 0.5 Å criteria.

### 3.3 Results on the protein-compound comparison set

The water position prediction performance and computation time of WatGNN were compared with GalaxyWater-CNN and 3D-RISM on the protein-compound comparison set, which has been used for performance comparison in a previous study^45^. The comparison in Figure 8 shows that RMSD between predicted positions and water positions from crystal structure is lowest for WatGNN, implying that WatGNN predicts closer to water positions in crystal structure than GalaxyWater-CNN and 3D-RISM. However, WatGNN showed worse average coverage than GalaxyWater-CNN. For example, WatGNN shows an average coverage with 1Å criteria of 49% and 76% when ***N***_pred_/***N***_cryst_= 1 and 3 respectively, while GalaxyWater-CNN shows coverage of 59% and 81%. The poor coverage could occur from the limitation of probe nodes that probe nodes are placed from non-carbon atoms, which is suitable for protein molecules that contain at least two non-carbon atoms per residue and can cover hydration sites of protein molecules with a small number of probe nodes but may not cover hydration site of ligand molecule that contains less amount of non-carbon atoms.

**Figure 8.**
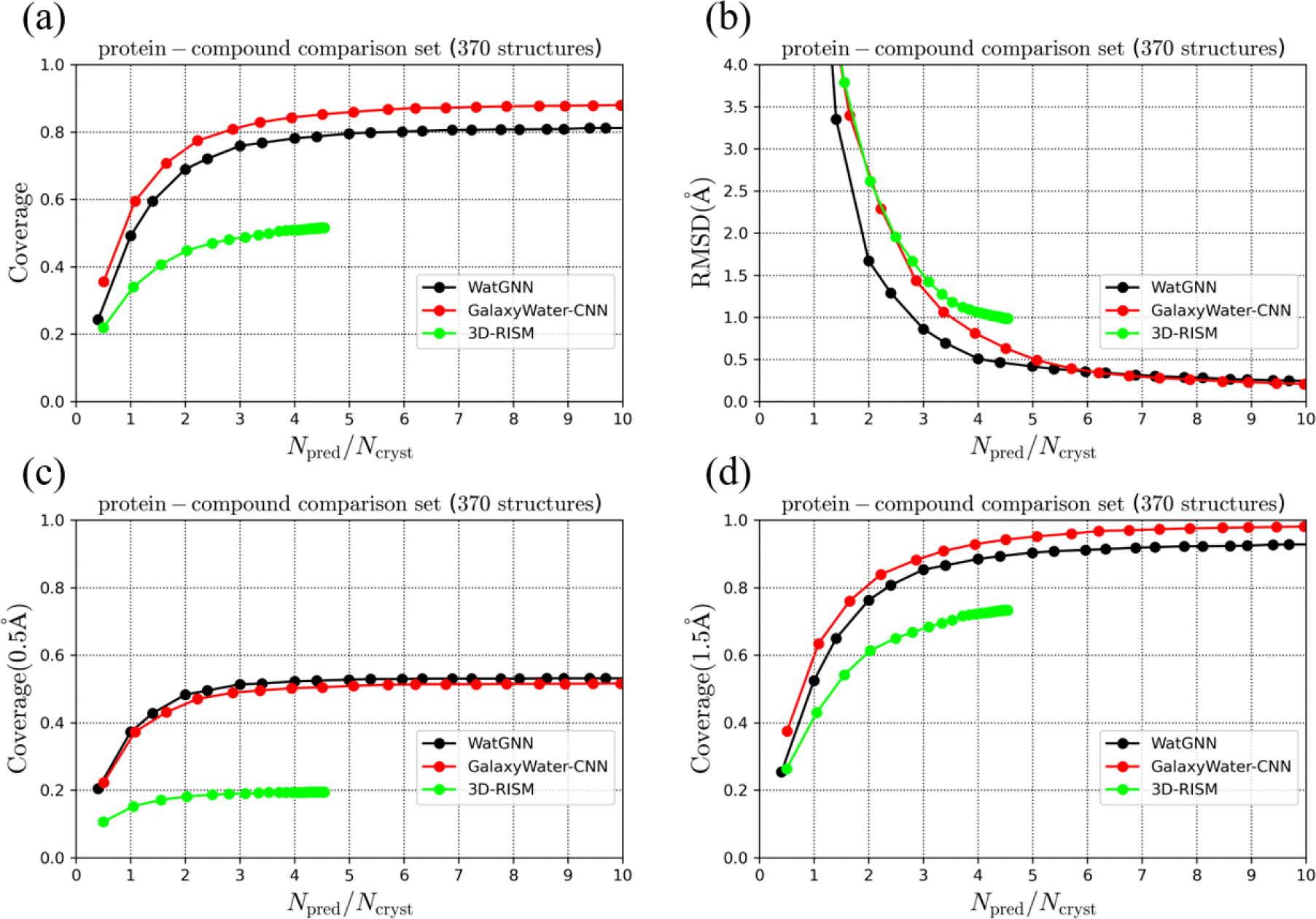
The prediction performance comparison of WatGNN on protein-compound comparison set: a), c), d) Average coverage, and d) Average RMSD of the predicted water molecules vs. the number of predicted water positions scaled by the number of water positions in the crystal structure was shown. For the average coverage, distance cutoff criteria of a) 1.0 Å, c) 0.5 Å, and d) 1.5 Å were used, respectively.

Figure 9 compares the water sites predicted with WatGNN and GalaxyWater-CNN as an example of prediction performance. The figure shows that predicted positions with WatGNN are closer to water sites in the crystal structure than those with GalaxyWater-CNN. Still, the prediction from WatGNN shows less coverage than the prediction from GalaxyWater- CNN.

**Figure 9.**
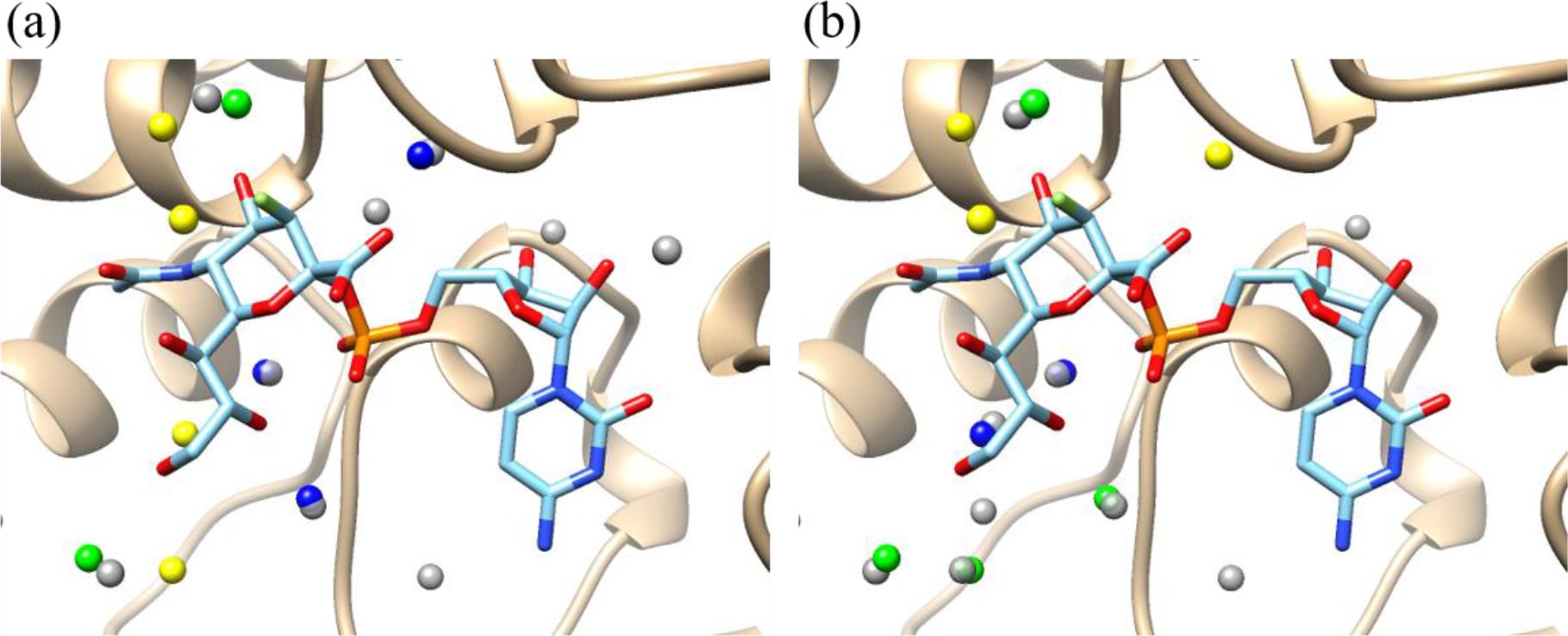
An example case of water site prediction (PDB ID: 2IHJ) done with (a) WatGNN and (b) GalaxyWater-CNN, with nine positions predicted, which is the number of crystallographic water molecules neighboring the ligand molecule. Gray spheres represent predicted water positions, and colored spheres (yellow, green, and blue) represent crystallographic water sites. Blue spheres indicate crystallographic water molecules with corresponding predicted water sites within 0.5 Å, and green spheres indicate crystallographic water molecules with corresponding predicted water sites within 1 Å, and yellow spheres indicate crystallographic water molecules for which no predicted water site is found within 1 Å. In this figure, water position prediction with WatGNN showed coverage of 56% (5 out of 9) for 1 Å criteria and coverage of 33% (3 out of 9) for 0.5 Å criteria. In comparison, GalaxyWater-CNN showed coverage of 67% (6 out of 9) for 1 Å criteria and 22% (2 out of 9) for 0.5 Å criteria.

## 4. Conclusions

WatGNN is a fast water position prediction method for protein structures that utilizes SE(3)-equivariant Graph Neural Network. Such a network enables fast prediction by considering each protein atom as a single node on a graph. In contrast, a Convolutional Neural Network considers a protein atom as data on multiple cells in the grid. Additionally, applying the constraint of roto-translational equivariance to the network ensures the independence of prediction results in rotation and translation of the input protein structure. To predict water positions with critical interaction with protein with low computational cost, a water site within 4.5 Å of the non-carbon atoms was predicted only, and probe nodes were introduced to consider protein atoms having multiple hydrogen bonds with water molecules. Also, with this approach, the prediction of water position can be made without using a grid, and the accuracy of predicted positions is not limited by grid spacing.

In this research, WatGNN showed more precise position prediction and less computation time than existing methods, such as GalaxyWater-CNN, GalaxyWater-wKGB, 3D-RISM, and FoldX. With fast and accurate water position prediction of WatGNN, critical interactions involving water molecules in protein structure could be considered with low computation time and less need for position refinement of predicted water positions.

Therefore, the effect of critical interaction involving water molecules could be considered for tasks that require low computation time for considering water effects, such as protein structures with structure sampling, protein-ligand docking, or designing biomolecules.

